# Quantifying the impact of dengue containment activities using high-resolution observational data

**DOI:** 10.1101/401653

**Authors:** Nabeel Abdur Rehman, Henrik Salje, Moritz U G Kraemer, Lakshminarayanan Subramanian, Simon Cauchemez, Umar Saif, Rumi Chunara

## Abstract

Dengue virus causes over 96 million cases worldwide per year and is expanding rapidly in geographic range, especially in urban areas. Containment activities are an essential part of reducing the public health burden caused by dengue, but systematic evidence on the comparative efficacy of activities from the field is lacking. To our knowledge, the effect of containment activities on local (sub-city) scale disease dynamics has never been systematically characterized using empirical containment and case data. We combine data from a comprehensive dengue containment monitoring system with confirmed dengue case data from the local government hospitals to estimate the efficacy of seven common containment activities in two urban areas in Pakistan. We use a modified version of the time series Suspected Infected Recovered framework to estimate how the reproductive number, *R*_0_, of the outbreak changed in relation to deployment of each containment activity. We also estimate the spatial dependence of cases based on deployment of each containment activity. Both analyses suggest that activities aimed at the adult phase of the mosquito lifecycle have the highest efficacy, with fogging having the largest quantifiable effect in reducing cases immediately after deployment. In examining the efficacy of containment activities contemporaneously deployed in the same locations, results here can guide recommendations for future deployment of resources during dengue outbreaks in urban settings.

## 1. Introduction

Dengue is a global threat; rapidly spreading with more than one half of the world’s population at risk for infection [1, 2]. Dengue virus is the most ubiquitous human arbovirus. It is transmitted primarily by *Aedes aegypti* mosquitoes, a vector which also transmits several other global threats including Zika, chikungunya and yellow fever [3]. Today, severe dengue is a leading cause of hospitalization and death among children and adults in urban areas in Asia, and Central and South America [4]. Dengue disproportionately affects urban areas in developing countries, which often have limited resources for containment and intervention activities [5, 6].

To date, the most common approach to reducing the burden of dengue is through prevention and containment of the vector population [7, 8]. Containment activities focused on vector control broadly fall into three categories: (i) activities targeted at reducing mosquito breeding sites (source reduction); (ii) activities targeted at the larval stage of the vector; and (iii) activities targeted at the adult stage of the vector [9]. While recent work has advanced efforts such as vaccines, genetically modified mosquitoes and Wolbachia-infected mosquitoes [10], these interventions are generally seen as a complement to containment activities [11], and may be prohibitively costly for many countries [12].

Despite the widespread use of containment activities, costing millions of dollars each year, the evidence base of how these activities reduce dengue risk is very limited. Existing research has largely focused on small controlled trials that estimate the effect of a containment activity by comparing treated and untreated populations [13, 14, 15, 16, 17]. Given the systematized nature of such studies, they generally focus on a small number of containment activities in a local, controlled environment; therefore the results may not be directly applicable to real-world settings, where external factors may impact the efficacy of the containment activities [18]. Further, nearly all efforts to quantify the effect of activities on vector control use markers of vector presence (e.g., household/container indices, Breteau indices) as the main outcome of measure, and do not incorporate disease incidence directly [19]. However, the link between vector measurements and dengue risk is poorly understood and a recent systematic review found little evidence of entomological indices such as the Breteau index being statistically associated with risks of dengue transmission [20, 21].

Here, we harness data from a novel containment monitoring system in two cities in Pakistan which has produced data on millions of instances of seven different types of containment activities, each linked with precise geolocation information. In parallel, there is detailed geo-location information on when and where dengue cases occurred in the cities. This provides a unique opportunity to estimate the impact of the different containment activities on the spatial distribution of cases, which we do using two statistical frameworks.

This study, as far as we are aware, considers the largest number of dengue containment activity types and instances alongside real field case data. Though the application and results are derived for dengue fever, this approach and findings can be informative for containment activity deployment for other arboviruses. Broadly, the results provide insight which can be used to help shape increasingly important decisions for resource allocation in Pakistan and other countries at risk of dengue and other vector-borne diseases.

## 2. Results

To quantify the impact of containment activities on disease incidence, we use data on 10,888 confirmed geocoded dengue cases reported in the cities of Rawalpindi (N=7,890 between January 1, 2014 and December 31, 2017, Fig. S3 and Fig. S5) and Lahore (N=2,998 between January 1, 2012 and December 31, 2017, Fig. S2 and Fig. S4). After a major dengue outbreak in 2011, the city of Lahore experienced two mild outbreaks in 2013 and 2016 while Rawalpindi has experienced outbreaks in each year since 2014. In addition, the date and precise location of 3,977,159 containment activities was recorded from the two locations (1,610,941 between January 1, 2014 and December 31, 2017 from Rawalpindi and 2,366,218 between January 1, 2012 and December 31, 2017 from Lahore) (Fig. S4, Fig. S5, Methods, Supplementary Text and Table S1).

### 2.1. Spatial Signature of Containment Activities

To understand the spatial effect of containment activities, we adapt an approach previously used to assess dengue spatial dependence at small spatial levels [22, 23]. The spatial dependence metric, *τ*, quantifies how the location and time of a case relates to the location and time of other cases. Specifically, *τ*_*i*_(*d*_1_, *d*_2_, *t*_1_, *t*_2_) is the relative probability of a case being reported in the distance window between *d*_1_ and *d*_2_, for cases *i*, within 30 days (*t*_2_ - *t*_1_, where *t*_1_ is the day when the case *i* developed first symptoms) compared to the expected probability of a case if there is no spatial dependence (the case clustering process is independent of space and time). Importantly, both the numerator and denominator of this metric are dependent on the spatiotemporal distribution of cases appearing in the same area and time-window, therefore controlling for exogenous heterogeneities that could create spatial or temporal clustering (e.g., variation in population density, hospital and healthcare use and reporting rates, and dengue seasonality). All details are explained in Methods and follow previous work [22].

We first calculate the spatial dependence between cases overall, and then specifically for cases in each of Rawalpindi and Lahore (Methods). Overall, when considering combined patients from both cities, we observe a 2.25 times (95% CI 2.16-2.33) increased probability of observing a case occurring within 50 m (*d*_1_=0 m and *d*_2_=100 m) radius and within 30 days of an index case, relative to the probability of a case occurring if clustering is independent in space and time, highlighting a strong spatial dependence between cases (Fig. 1). This falls to 1.37 (95% CI 1.33-1.40) at a distance of 1.25 km (*d*_1_=1 km and *d*_2_=1.5 km) and 1.0 (95% CI 0.98-1.02) at a distance of 4.55 km (*d*_1_=4.3 km and *d*_2_=4.8 km). When calculating spatial dependence separately for cases in each city, we observed a 2.21 times (95% CI 2.14-2.28) and 1.46 times (95% CI 1.29-1.59) increased probability of observing a case occurring within 50 m (*d*_1_=0 m and *d*_2_=100 m) radius and within 30 days of an index case (Fig. S6) in Rawalpindi and Lahore, respectively. The lower level of spatial dependence in Lahore, as compared to Rawalpindi, suggests variation in spatial dependence of cases, across different locations and times, should be accounted for when studying the effect of containment activities.

**Figure 1:**
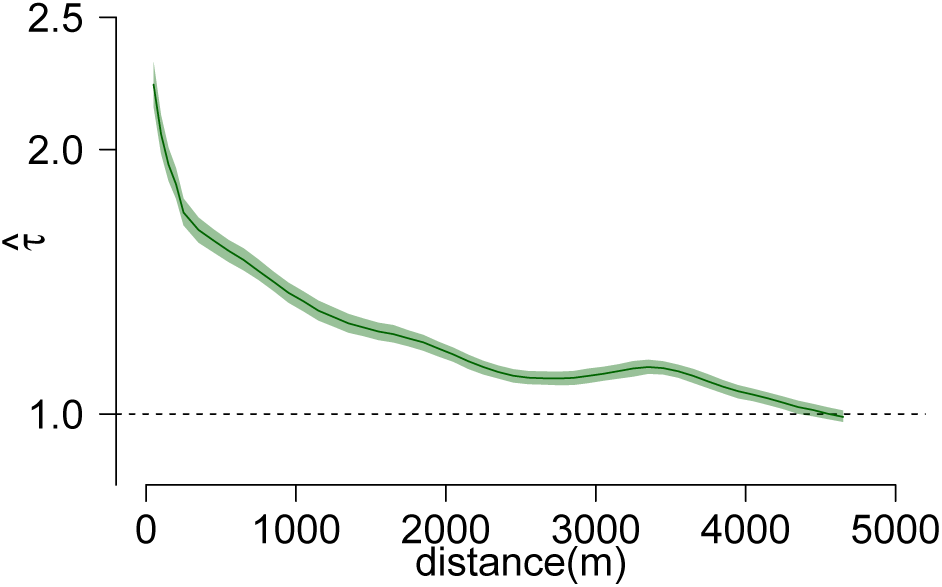
Spatial dependence of cases occurring within 30 days (cases from Lahore and Rawalpindi). The spatial window of the analysis (*d*_2_ – *d*_1_) is maintained at 500 m when *d*_2_ is greater than 500 m, and observations are made by sliding the window at intervals of 100 m. For *d*_2_ less than 500 m, *d*_1_ is equal to zero and observations are made by increasing *d*_2_ at intervals of 100 m. Spatial dependence estimates are plotted at midpoint of the spatial window. The time window *t*_2_ – *t*_1_ is set to 30 days. 95% CI from bootstrapping 100 replications is shown as green shaded area around estimate.

We then study the result of different containment activities on the spatial dependence between cases. Of the 9,268 geo-tagged cases in Rawalpindi and Lahore between 2014 and 2017, 531 were assigned IRS, followed by larviciding (*n*=275) and fogging (*n*=162) (Table S2). A total of 742 cases had multiple containment activities in their spatio-temporal proximity and hence were not used as index cases in the study. As underlying spatial dependence may differ by different areas in the city or at different times during an epidemic season, for each case where a containment activity was performed, we identify a matched control where no activity occurred. Matched-controls occurred within 30 days and 1000 m of the containment-case but which were not in immediate vicinity of any containment activities. We define *ξ*_*a*_(*d*_1_, *d*_2_), as the ratio of the spatial dependence in distance window *d*_1_ and *d*_2_, as measured through *τ*, for cases which were in proximity of containment activity *a*, to the same measure for the matched control. Values of *ξ*_*a*_ below 1 signify that the relative probability of new cases appearing around a case which was in proximity of a containment activity is lower compared to that of a control case, after adjusting for underlying clustering in space and time, which is consistent with a positive impact from the containment activity. Values of *ξ*_*a*_ around 1 indicate no impact of the activity.

We calculate the *ξ*_*a*_ values for each containment activity, *a*, using combined data from both cities and for each city separately (Fig. 2, Fig. S7 and Fig. S8). When considering combined data, we find a consistent reduction in probability of new dengue cases in proximity of indoor residual spray (IRS) and fogging (Fig. 2). There was a 0.9 reduced probability of a case occurring within 50 m (*d*_1_=0 m and *d*_2_=100 m) and in the next 30 days of cases for which IRS occurred immediately after and in the immediate vicinity (95% CI: 0.81-0.99) (details in Methods). For fogging, this value was 0.80 (95% CI: 0.66-0.96). By 750 m (*d*_1_=500 m and *d*_2_=1000 m) for IRS and 1050 m (*d*_1_=800 m and *d*_2_=1300 m) for fogging, there was no difference (*ξ*_*a*_=0.99) in probability of new cases around the containment cases and the controls (Table S3). In contrast to fogging and IRS, there was no consistent reduction in probability of new cases in proximity of any other containment activity (Fig. 2). This lack of effect is most clearly visible for larviciding which had the most number of cases amongst activities which had no effect (*n*=275). Due to the low number of cases in proximity of tap fixing (*n*=25), the resulting plot for this activity indicate structural uncertainty and are not interpretable. Findings were consistent when we varied the maximum distance of matched controls (Fig. S9) and when considering cities separately (Fig. S7, Fig. S8 and Table S2).

**Figure 2:**
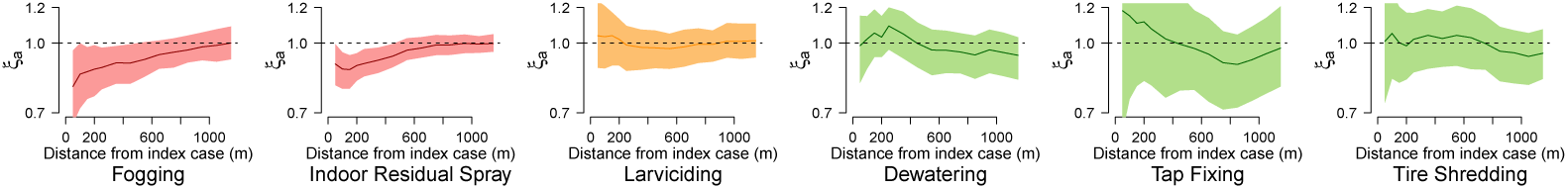
Variation in the effect of containment activity, *ξ*_*act*_, versus the distance (in meters) from index cases using combined data from Rawalpindi and Lahore. Values of *ξ*_*act*_ are calculated using control and containment cases which appear in an *m*=1000 m radius of each other. The spatial window of the analysis (*d*_2_ – *d*_1_) is maintained at 500 m when *d*_2_ is greater than 500 m, and observations are made by sliding the window at intervals of 100 m. For *d*_2_ less than 500 m, *d*_1_ is equal to zero and observations are made by increasing *d*_2_ at intervals of 100 m. Spatial dependence estimates are plotted at midpoint of the spatial window. Values below 1 show a lower probability of new cases appearing around a case in proximity of a containment activity, compared to a control case. The time window *t*_2_ – *t*_1_ is set to 30 days. 95% CI from bootstrapping 100 replications are shown as shaded areas around estimates. Activities targeted at adult stage of mosquito are shaded red, activities targeted at larval stage shaded orange, and activities targeted at source reduction are shaded green.

### 2.2. *Impact of Containment Activities on* R_0_

To understand the effect of containment activities on the transmission potential of the outbreak and cases over time, we fit a Time Series Susceptible Infected Recovered (TSIR) model for sub-city spatial units from both cities (Methods) using the adjusted reported cases. Additionally, we create separate TSIR models for each city (Supplementary Text).

This modeling approach is useful as it allows us to account for environmental drivers, which are very pertinent in dengue epidemiology, and it assesses transmission potential through a standardized metric, *R*_0_. In both Lahore and Rawalpindi, we observe high dengue activity during the post monsoon months, September-November, which highlights the importance of climate in the reproduction of dengue vector (Fig. S2 and Fig. S3). Given that nearly half of dengue cases are asymptomatic and given that our dataset primarily comprises of data from public hospitals, we adjust the reported cases for under-reporting (Methods and Supplementary Text) [1]. We also assessed sensitivity of results based on this reporting rate; showing no changes in the overall results (Fig. S14).

Each city is divided into spatial units (*N*=10 for Lahore and *N*=14 for Rawalpindi), based on administrative boundaries to model localized dengue transmission. We included containment activities, environmental data (temperature and rainfall), and population density as part of the model to identify the effect of each of these parameters. Appropriate delays, to account for vector life cycle and transmission of virus from vector to human were added, and the residual effect of containment activities was accounted for, to model realistic transmission of dengue accurately infer the effect of each parameter (Supplementary Text). To access the utility of containment data, we train additional variants of the TSIR model using only environmental parameters and population density.

The model trained on data from spatial units from both cities, using only environmental parameters and population density, provided a good fit (adjusted *R*^2^ = 0.63), and the addition of containment activities to the model improved the fit (adjusted *R*^2^ = 0.65). For the model trained only on data from spatial units in Rawalpindi, the addition of containment activities improved the adjusted *R*^2^ from 0.78 to 0.81. Similarly, for Lahore the model incorporating containment activities improved the adjusted *R*^2^ from 0.73 to 0.76 (Akaike information criterion (AIC) values also reported in Table S7). Overall, for the model trained on combined data, the reproductive number was 2.82 (at mean temperature and precipitation values; 25.5 Celsius and rainfall for 2 days during a 2 week period), if all containment activity coefficients are set to zero. For Lahore the *R*_0_ was 1.59 (at 26 Celsius and 2 days of rainfall), and for Rawalpindi the *R*_0_ was 1.79 (at 24.9 Celsius and 2 days of rainfall).

Our results illustrate varied relationships between an increase in the amount of containment activities and cases over time, for each activity as it was deployed in Lahore and Rawalpindi, and using *R*_0_ (Fig. 3, Fig. S12 and Fig. S13). We quantify the amount of a containment activity in instances, where an instance during a single time-step (2 weeks in our study) represents the sum of the number of activities performed during the time-step, and the residual effect of any activities performed in previous weeks (Supplementary Text). For example for fogging, which has no residual effect, an instance at time *t* represents only the number of activities performed in a spatial unit at *t*. In contrast, for IRS which has a residual effect, instances at time *t* represent the sum of the number of IRS activities performed at time *t* and the residual effect of IRS activities performed in the previous six time-steps (the residual effect of IRS is three months).

**Figure 3:**
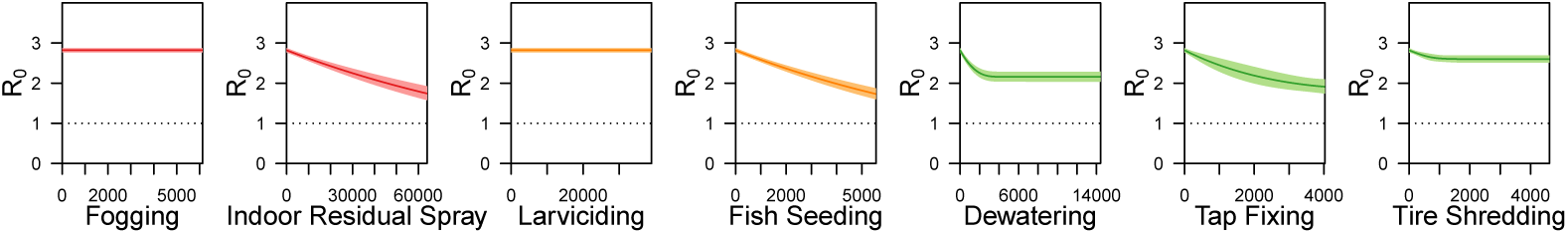
Variation in reproductive number (*R*_0_) of dengue, with variation in instances of containment activity, estimated from the model trained using data from (*N*=10 spatial units) in Lahore between 2012 and 2017, and (*N*=14 spatial units) in Rawalpindi between 2014 and 2017. X-axis represents the total number of containment activities performed, in a spatial unit, in a lagged time step and any residual effect from previous weeks.

Of the adulticides, we find an increase in IRS to be related to a decrease in *R*_0_ of dengue in both Lahore and Rawalpindi, as well as when data from both cities is modelled as part of a single model. Specifically, additional deployment of approximately 4,800 IRS activities in a spatial unit was related to a 0.1 decrease in the *R*_0_ of dengue. In contrast, fogging was related to a decrease in the *R*_0_ of dengue only in Lahore. Among containment activities targeted at the larval stage of mosquitoes, larviciding showed no effect on *R*_0_ in either city or when data from both cities was trained together, while fish seeding was only related to a decrease in *R*_0_ when data from both cities was trained in a single model. Among source reduction activities, tap fixing was related to a decrease in *R*_0_ in Lahore and in the model with combined data from both cities. Tire shredding was related to a decrease in *R*_0_ in Rawalpindi, and when analyzing combined data from both cities, but the effect of this activity was not statistically significant in Rawalpindi. Dewatering was only related to a decrease in *R*_0_ when data from both cities was trained in a single model. Results across all models are summarized in Table S4.

## 3. Discussion

Data from the dengue containment activity monitoring system deployed in the Punjab province, Pakistan in 2012 was used; which, to our knowledge, monitors the largest number and types of containment activities. The system captured millions of containment activity events over a seven-year period (Table S1), each event linked to precise geo-coordinates. Combined with geo-location of patients, this allowed us to systematically examine the effect of multiple containment activities on sub-city scale disease dynamics, which has never before been characterized using empirical activity and case data.

We examined the relationship between deployed instances of each containment activity type and the spatial dependence of geo-located dengue cases in their proximity, in the cities of Rawalpindi and Lahore between 2014 and 2017. This method allows generation of unbiased estimates in the midst of exogeneous heterogeneities that could create spatial or temporal clustering (e.g., variation in population density, hospital and healthcare use and reporting rates, and dengue seasonality). The result is quantification of both the maximum reduction in dengue transmission in the vicinity of a particular type of activity, as well as the maximum distance at which this reduction in dengue transmission is evident. Notably, the method and results provides novel empirical results insights into the comparative efficacy of fogging and indoor residual spray using real case and containment activity data.

The time series modelling of dengue cases in Lahore and Rawalpindi enabled us to assess the relation between the *R*_0_ of dengue and amount of containment activities, as deployed. Results from this approach are based on empirical field data, consider multiple interventions and use a precise and standardized measure of efficacy (*R*_0_) in contrast to studies based on simulated data and models, or using proxy measures for dengue transmission [19]. The results show that training a separate model for spatial units in each city provides a better fit to data and hence results from models trained for individual cities get precedence over the model trained on combined data.

The spatial dependence of dengue cases reported here is consistent with that reported in previous work using dengue case data from Bangkok. The spatial dependence at 200 m, presented in [22] is 1.82 (95% CI: 1.45-2.16) is comparable to 1.87 (95% CI: 1.81-1.93) observed in the two cities in Pakistan in our study. Further, the values of 1.83 and 1.45 observed in Rawalpindi and Lahore respectively also lies within the confidence interval. Results from the spatial signature analysis show that application of IRS and fogging spray, in the vicinity of a dengue case, result in reduction of the generation of new cases by 10% and 20% respectively. Additionally, IRS and fogging are shown to be effective (*ξ*_*a*_ below 1) up to a distance of 750 m and 1050 m respectively. Similar trends are observed based on the results of time series modelling of containment activities. Increases in IRS and fogging are related to decreases in the reproductive number of dengue in Lahore, though results from Rawalpindi specific model only show a statistically significant effect from IRS. This could be due to the fact that TSIR models assume that activities and cases are uniformly distributed in each spatial unit considered. If the assumption is violated and activities are not performed in the direct vicinity of cases, then the resulting effect from the model may not be completely accurate [24].

Results from both the spatial dependence method and timeseries modelling did not find larviciding to be effective. These results are consistent with a recent systematic review, which found Temephos (a chemical used in larviciding) to be only effective in reducing entomological indicators, but found no evidence of its association with reduction in disease transmission. At the same time, the results highlight that while containment activities can be effective under laboratory conditions, the effectiveness does not translate exactly in the field in reducing dengue transmission. This signifies the utility of studies such as this which examine effectiveness of containment activities using real case data. For example, there is conflicting evidence regarding the effectiveness of fish seeding in the literature [13, 25]. Our time series method did not find fish seeding to be effective in either city, and due to a minimal number of cases which were adjacent to only fish seeding activities, no inference about the effectiveness of fish seeding could be made from the spatial dependence method.

Among source reduction containment activities, we find no activity to be effective using the spatial dependence method. Using the TSIR model, we find an increase in tap fixing in Lahore and increase in dewatering in Rawalpindi to be associated with a decrease in the reproductive number of dengue.

Quantitatively, our results corroborate existing knowledge about the role of rainfall and temperature in dengue transmission by showing increases in *R*_0_ with increases in temperature and number of rainfall days [26, 27] (Supplementary Text). We also find an increase in population density is related to an increase in *R*_0_, when considering data from both cities separately (Supplementary Text).

It should be noted that results from this study are only relevant to the spatial dependence of cases or relationship between containment activity deployment and *R*_0_ after dengue cases have started to appear. Results from the study do not explain the effect of a containment activity on the overall dengue burden, or on delaying or preventing the appearance of first cases. A separate, and longitudinal analysis would be required to evaluate the preventive effectiveness of each containment activity. As well, as with any study based on human reported data, there could be a chance of sampling bias in the containment activity reports. Such a bias would have to have a systematic spatial or temporal dependence in order to impact results; thus we deem the assumption that such a bias would not affect the results fair. Further, while we consistently observe a short-term positive impact of IRS on dengue incidence, we were unable to assess the longer-term impact of the containment activities and we cannot rule out these containment activities simply delay infection to future time points [28].

In conclusion, results of this study regarding the relationship of different containment measures with the spatial dependence of dengue cases or the *R*_0_, provide specific insight regarding dengue in urban settings. More broadly, these results and the models and methods used to derive them – are relevant to a growing number of global health concerns related to the *Aedes aegypti* mosquito, including the Zika virus and chikungunya, which are also known to particularly impact urban areas. Further, the methods presented in the work lay groundwork for future studies aimed at studying the effect of containment from observational data collected from the field.

## 4. Methods

### 4.1. Containment Activities Data

Modern technology was applied by the Punjab Information Technology Board to track containment activities carried out by the Punjab Health Department. Mobile phones were distributed to health care workers to record their activities since 2012 using a mobile application (Supplementary Text and Fig. S1). Government workers were asked to take a picture before and after performing the containment activity as a verifiable proof that the activity had been performed (Supplementary Text). Global positioning system (GPS) coordinates of the location, time stamp, and pictures of the performed activity were automatically submitted to a centralized server where they were monitored. Data on dengue containment activities for the period January 1,2012 to December 31, 2017 was received. This consisted of 7,281,932 containment records, each including the name of the containment activity, a time stamp of when the activity was performed and the GPS coordinates for the location of where it was performed. After excluding those activities performed outside the boundaries of the two cities, we were left with a total of 2,366,218 containment activity instances in Lahore between January 1, 2012 and December 31, 2017, and 1,610,941 activity instances performed in Rawalpindi between January 1, 2014 and December 31, 2017. For the TSIR model, we used the GPS coordinates to map each containment activity data point to a spatial unit.

### 4.2. Epidemiological data

Data regarding confirmed dengue cases, for the same time period as the containment activities, was retrieved from the Government of Punjabs centralized patient portal system. Precisely geo-tagged information linked to each case was available starting in 2014 (spatial unit level data was available from 2012-2014 for Lahore) (Supplementary Text). A total of 2,998 cases were reported in Lahore between January 1, 2012 and December 31, 2017. In Rawalpindi a total of 7,890 confirmed dengue cases were reported and geo-tagged between January 1, 2014 and December 31, 2017.

### 4.3. Environmental Data

City-wide daily mean temperature and mean precipitation estimates, for both cities, were obtained from the Pakistan Meteorological Department for time series method (www.pmd.gov.pk accessed August 27, 2018). As previously shown these climate factors directly affect mosquito survival, reproduction, and development and thus their abundance.

### 4.4. Spatial Dependence of Cases

First, to characterize the spatial dependence of cases we compute the probability of a case occurring between times *t*_1_ and *t*_2_, and within distance range *d*_1_ and *d*_2_ of a given case versus the expected probability if the clustering processes were independent in space and time:

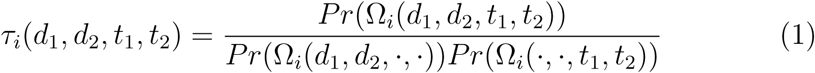

where Ω_*i*_(*d*_1_, *d*_2_, *t*_1_, *t*_2_) is the set of cases between *d*_1_ and *d*_2_ (in meters) and temporal window of *t*_1_ and *t*_2_ (in days) of case *i*; Ω_*i*_(*·, ·, t*_1_, *t*_2_) is the set of cases in temporal window *t*_1_ to *t*_2_ of case *i* independent of space, and Ω_*i*_(*d*_1_, *d*_2_, *·, ·*) the set of cases within spatial window *d*_1_ and *d*_2_ of case *i*, independent of time. For our analysis, we use a fixed time window of 30 days: *t*_1_ is selected as the day when the patient experienced first symptoms of dengue virus, and *t*_2_ = *t*_1_ + 30. This time window is chosen to ensure that cases considered are from the same transmission chain, though we perform sensitivity analysis using additional time windows (Fig. S10). Dependence is then observed across variation in the distance window.

Then, the overall spatial dependence of new cases appearing around cases labelled *s* (labelling is defined in the next subsection) is estimated as:

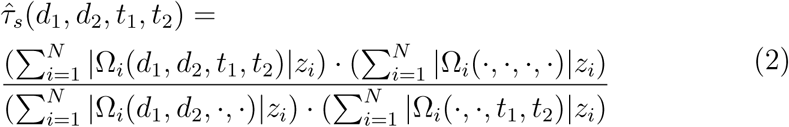

where *z*_*i*_ is 1 if the case is labelled *s, N* is the total number of cases in the dataset regardless of their label, and Ω_*i*_(·,·,·,·) is the set of all cases in the dataset.

### 4.5. Spatial Signature of Containment Activities

To identify the impact of containment activities on the spatial dependence of dengue cases (the “spatial signature” of an activity) we first label all cases in the dataset as either a “containment” or a “control”. A case is labelled as *s* = *a* if only the containment *a* was performed in a 20 meter radius and time window of the past 30 days of the case before the first symptom appeared. Only cases for which a single containment activity was performed in the surrounding area are included in the analysis, to ensure only the effect of a single type of containment activity is being measured. A case is labelled a control, *s* = *c*, if no containment activity was performed in a 20 meter radius and time window of the past 30 days of the case before the first symptom appeared. The *tau* metric measures clustering dynamics, however there are factors such as population variation, reporting biases and availability of vegetation and water for growth of vector, can also play a role in variation of the number of cases that would be expected in a given location and time. Thus, to compare clustering while controlling for such factors, we compare clustering around cases that have a similar epidemiological context. For a given set of containment cases labelled *a*, we select a subset of cases, *a*^*’*^, such that each case in *a*^*’*^ has a matching control case. A matching control case is defined as a control case which is within a radius of *m* meters, and was reported within 30 days of the containment case. We assess how values of *m* of 500, 1,000 and 2,000 (Fig. 2 and Fig. S9) impact the results. For each containment case *a*^*′*^, we randomly select a matching control case and represent the set of matching control cases as *c*^*′*^_*a*_. The spatial signature of containment activity *a, ξ*_*a*_, is then calculated as:

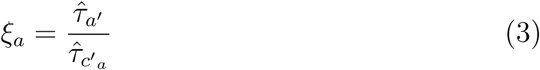

### 4.6. Impact of Containment Activities on R_0_

We model the incidence of dengue using a time-series susceptibleinfectedrecovered (TSIR) model of viral incidence previously used to reconstruct dengue dynamics in Asia (Supplementary Text) [29, 30]. The city of Lahore is divided in (*n*=10) and the city of Rawalpindi in (*n*=14) spatial units, and localized transmission of dengue is modelled at each spatial unit. The reported cases, in each spatial unit, are first reconstructed to account for under-reporting. The reported number of cases, 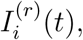 are first smoothed, then multiplied with the inverse of the reporting rate *rr*, and the product is used as the mean of Poisson distribution (Supplementary Text, and Table S5 and S6). The number of infected individuals, *I*_*i*_(*t*), are selected at each time step from the distribution. This reconstruction methodology, used in previous infectious disease modeling work [31], gives the advantage of capturing tails of the epidemic curve in a realistic, continuous manner. Our model incorporates environmental parameters in the transmission rate to account for variation in vector population density. We use two weeks as the time step in our study, consistent with the generation interval and previous studies which model the transmission of dengue [32, 30] (Supplementary Text). The general TSIR model is defined via the following equations:

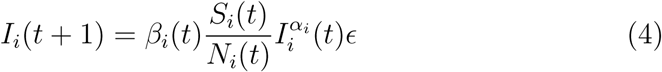

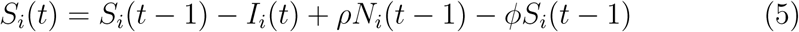

where *I*_*i*_(*t*), *S*_*i*_(*t*) and *N*_*i*_(*t*) are the infected, susceptible and total population during time step *t* in spatial unit *i, ρ* is the bi-weekly birth rate, *φ* is the biweekly death rate, *α*_*i*_ is the mixing coefficient in spatial unit *i*, and *β*_*i*_(*t*) is the transmission coefficient during time step *t*. The error term *E* is assumed to be an independent and identically log-normally distributed random variable.

We endogenize containment activities in the transmission coefficient *β*_*i*_(*t*). This decision reflects the fact that containment activities reduce the contact rate between humans and mosquitoes, which results in a reduction of the transmission rates from human to mosquito to human [33]. The transmission coefficient *β* for equation 4 is parameterized as:

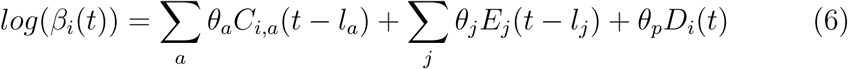

where *l*_*a*_ and *l*_*j*_ are time steps containment activities *a* and environmental parameters *j* were lagged respectively (Supplementary Text). *C*_*i,a*_(*t-l*_*a*_) is the number of times per squared kilometer containment activity *a* was performed in spatial unit *i* during week (*t - l*_*a*_). *E*_*j*_(*t - l*_*j*_) is the value of environmental parameter *j* during week (*t-l*_*j*_). *D*_*i*_(*t*) is the population density in spatial unit *i*. The residual effect of each containment activity is added based on existing knowledge (see section Transmission cycle of dengue and timing and residual effect of containment activities in Supplementary Text).

To calculate the value of *β*_*i*_(*t*), the value of *β* for each town at each time step, a single model is used to find the best fit for parameters: *θ*_*a*_, *θ*_*j*_, *θ*_*p*_, based on the number of each containment activity and environmental parameters as well as all non-zero cases data point in each town, *i*, at every time step (equation 6). We use Shape constrained additive model (SCAM) to fit this relationship. Shape constrained additive models are an extension of generalized additive models (GAMs) which provide the advantage of using existing knowledge about the relationship of the response variable with the explanatory variables [34, 35]. This prevents noise from being included in the shape of splines from the GAM. Containment activities are modeled as monotonically decreasing splines while environmental parameters and population density are modeled as monotonically increasing splines. The smoothing parameters are estimated using maximum likelihood. Finally, using estimates of *θ*_*a*_, *θ*_*j*_, and *θ*_*p*_ from the SCAM model and equation 6 and 7, we identify the variation in *R*_0_ (reproductive number of dengue) by variation in the amount of each containment activity. The *R*_0_ is calculated by the following equation:

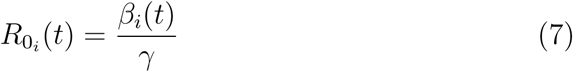

where, *γ* is the recovery rate and is equal to 1 time step in our study, given the fact that infected patients are immediately admitted in the hospital and removed from the infected population. The reproductive number can be defined as the number of secondary infections a primary infection can cause over the course of its infectious period [36]. If *R*_0_ is greater than 1, then the disease will spread exponentially, while an *R*_0_ below 1 means that the disease will not spread.

